# Four new induced pluripotent stem cell lines produced from northern white rhinoceros with non-integrating reprogramming factors

**DOI:** 10.1101/202499

**Authors:** Marisa L. Korody, Cullen Pivaroff, Thomas D. Nguyen, Suzanne E. Peterson, Oliver A. Ryder, Jeanne F. Loring

**Author notes:** These authors contributed equally to this work (Marisa L. Korody, Cullen Pivaroff). Current address: Genea BioCells USA Inc. La Jolla, CA 92037 USA.

## Abstract

Since we first published methods for generation of induced pluripotent stem cells from endangered species^1^, we have developed improved non-integrating methods for reprogramming the functionally extinct northern white rhinoceros, and generated iPSCs from four more individuals. Our work is part of a long-term plan for assisted reproduction for conservation of endangered species.

## Introduction, Results, and Conclusions

Species are disappearing from the earth at an alarming rate, and a novel approach that may help in their survival is generation of induced pluripotent stem cells (iPSCs) from highly endangered species. iPSCs can divide indefinitely in culture and be differentiated into any cell type in the body, including eggs and sperm. Thus, it may be possible to use assisted reproduction methods to generate new individuals of a species. Toward this end, in 2011 we reported reprogramming of two endangered species, the drill (*Mandrillus leucophaeus*) and the northern white rhinoceros (*Ceratotherium simum cottoni*: NWR)^1^, using integrating lentivirus containing the four human reprogramming factors (POU5F1, SOX2, MYC, KLF4)

The NWR is functionally extinct, with only three non-reproductive individuals remaining alive, which makes it a worthy candidate for following through on the innovative strategy to use iPSCs in their conservation. Here we report fundamental improvements in reprogramming methods for the NWR, and the generation of iPSCs from four more NWR individuals, bring the total to five reprogrammed individuals. We developed species-specific culture conditions and optimized methods for delivering the human reprogramming factors. We report for the first time the efficient production of NWR iPSCs using the non-integrating Sendai virus (CytoTune™ 2.0 ThermoFisher), which is greatly preferable to integrating methods for use of the iPSCs for generation of gametes for assisted reproduction. We performed in-depth analysis of iPSCs from “Angalifu,” the last NWR male to live in the United States, residing at the San Diego Zoo Safari Park from 1990 until his death of natural causes in 2014.

Fibroblast cell lines were obtained from the San Diego Zoo’s Frozen Zoo®^2^ repository of biomaterials, which contains cell lines from 12 NWR individuals and captures a high level of genetic diversity within the species. These cell lines were established opportunistically from skin biopsies and were cryopreserved between 1979 and 2016. In order to improve generation of iPSCs from these fibroblasts, we optimized many aspects of the protocol, including culture
medium formulation, cell density, viral titer, and timing of transfection (see Supplemental Methods).

The new NWR iPSC colonies have a distinctive morphology that we noted in our earlier publication^1^ (Figure 1a). Immunocytochemistry (ICC) showed that the iPSCs were positive for canonical markers of pluripotency [OCT4 (also known as POU5F1), SOX2 and NANOG] (Figure 1b) and the iPSCs differentiated via embryoid body (EB) formation expressed markers for each of the germ layers (Figure 1c)^1, 3^. qPCR analyses of the iPSCs confirmed expression of pluripotency markers (Figure 1d). Interestingly, the NWR iPSCs expressed low levels of KLF4 compared to the source fibroblasts, which is consistent with the observed lack of insertion of KLF4 lentivirus in our initial iPSC derivation^1^. The karyotype of Angalifu’s iPSC line matches that of the initial fibroblast cell line and is normal (2n=82: Figure 1e).

**Figure 1:**
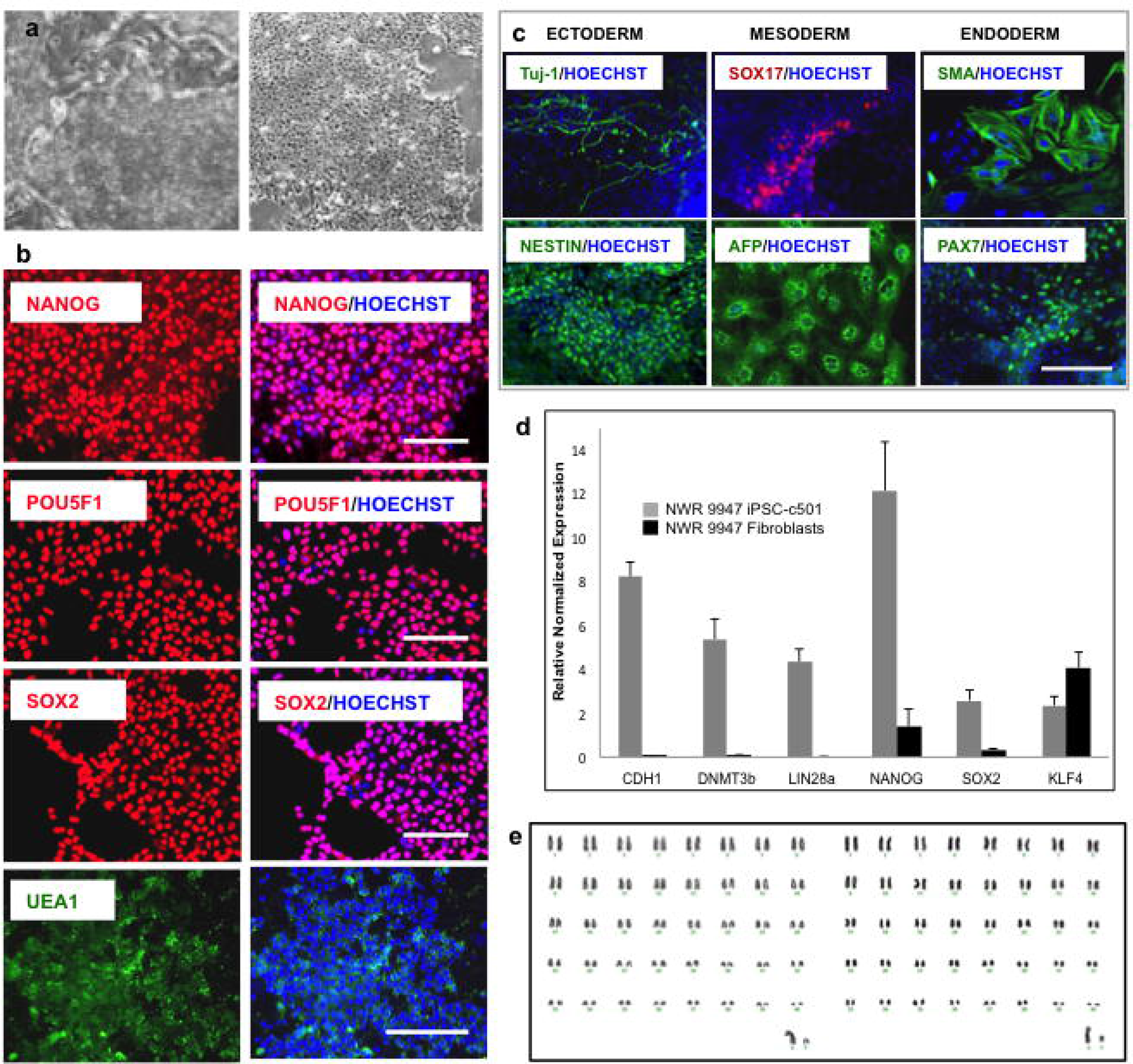
Characterization of northern white rhinoceros iPSCs. **a**) Phase contrast micrographs of feeder-dependent NWR iPSCs (left) and feeder-free NWR iPSCs (right). **b**) Immunocytochemistry for nuclear pluripotency markers NANOG, OCT4/POU5F1, SOX2, and cell surface marker UEA1. **c**) Immunocytochemistry for germ layer markers in differentiated embryoid bodies. Ectoderm: Tuj1 (TUBB3) and Nestin; Mesoderm: SOX17 and AFP (Alpha Fetoprotein); Endoderm: SMA (Smooth Muscle Actin) and PAX7. **d**) RTPCR of pluripotency markers using rhino-specific primers. **e**) Karyotyping confirmed that there were no chromosomal differences between the source fibroblasts (left) and the iPSCs (right).

These NWR iPSCs offer hope for rescue of this nearly extinct species and are part of a larger plan to restoration of this species through *in vitro* differentiation of functional germ cells and assisted reproduction ^4^. The reprogramming and characterization methods reported here should serve as a template for reprogramming and possible rescue of additional endangered species.

## References

1. Ben-Nun, I.F. et al. Induced pluripotent stem cells from highly endangered species. Nature Methods 8, 829–U889 (2011).

2. Chemnick, L.G., Houck, M.L. & Ryder, O.A. in Conservation Genetics in the Age of Genomics. (eds. G. Amato, R. DeSalle, H.C. Rosenblum & O.A. Ryder) 124–130 (Columbia University Press, New York; 2009).

3. Wang, Y.-C. et al. Specific lectin biomarkers for isolation of human pluripotent stem cells identified through array-based glycomic analysis. Cell Research 21, 1551–1563 (2011).

4. Saragusty, J. et al. Rewinding the Process of Mammalian Extinction. Zoo Biology 34, 280–292 (2016).

